# GenoDrawing: An autoencoder framework for image prediction from SNP markers

**DOI:** 10.1101/2023.03.06.531351

**Authors:** Federico Jurado-Ruiz, David Rousseau, Juan A. Botía, Maria José Aranzana

## Abstract

Advancements in genome sequencing have facilitated whole genome characterization of numerous plant species, providing an abundance of genotypic data for genomic analysis. Genomic selection and neural networks, particularly deep learning, have been developed to predict complex traits from dense genotypic data. Autoencoders, a neural network model to extract features from images in an unsupervised manner, has proven to be useful for plant phenotyping. This study introduces an autoencoder framework, GenoDrawing, for predicting and retrieving apple images from a low-depth single nucleotide polymorphism (SNP) array, potentially useful in predicting traits that are difficult to define. GenoDrawing demonstrated proficiency in its task while using a small dataset of shape-related SNPs, and multiple experiments were conducted to evaluate the impact of SNP selection and shape relation. Results indicated that the correct relationship of SNPs with visual traits had a significant impact on the generated images, consistent with biological interpretation. While using significant SNPs is crucial, incorporating additional, unrelated SNPs results in performance degradation for simple NN architectures that cannot easily identify the most important inputs. The proposed GenoDrawing method is a practical framework for exploring genomic prediction in fruit tree phenotyping, particularly beneficial for small to medium breeding companies to predict economically significant heritable traits. Although GenoDrawing has limitations, it sets the groundwork for future research in image prediction from genomic markers. Future studies should focus on using stronger models for image reproduction, SNP information extraction, and improved dataset balance in terms of shape for more precise outcomes.

## 1 Introduction

Advances in high throughput genome sequencing and genotyping methods have brought to reality the whole genome characterization of many plant species, including models and crops, as well as the analysis of diversity at population level. Under this scenario of huge amounts of genotypic data, the link between genotype and phenotype in crops has the tremendous potential to identify genes or genome regions involved in the natural variation of relevant agricultural traits, as well as to predict the performance of offsprings in specific environments. For traits controlled by major genes or moderate to strong Quantitative Trait Loci (QTLs), linkage mapping in biparental families or genome-wide association analysis (GWAS) in germplasm collections allow the development of markers that can, eventually, be used for marker-assisted breeding (MAB). By contrast, complex quantitative traits regulated by multiple QTLs with minor effects are difficult to predict with few markers. Genomic selection (GS) models have been developed to overcome this limitation [3, 34, 32]. In short, they predict complex traits from dense genotypic data and a set of qualitative, continuous, or discrete measurable trait descriptors in a training population to ultimately estimate the performance of the individuals of a breeding population from their genomic profile. More recently, Neural Networks (NN) are being suggested as a powerful tool for genomic prediction that may surpass some challenges associated to the classical GS models, such as the requirement of assuming data distributions of the linear models, or the requirement of priors’ specifications in Bayesian models [21]. Neural networks and deep learning (DL) are subfields of artificial intelligence that use multiple interconnected nodes, called artificial neurons, to process information and make predictions. These networks can have multiple layers, allowing for complex and non-linear analysis of data. Deep learning algorithms have demonstrated improved accuracy and speed compared to other artificial intelligence-based methods [29]. Deep neural networks (DNN) have been already applied for genomic prediction in several fields, using real [18, 28] and simulated [2] SNP marker data, stimulated by the growing availability of easy-to-use DNN frameworks such as PyTorch [20] and TensorFlow [1]. Within the DL fields of study, generative models (i.e. unsupervised models that learn from patterns to finally generate new examples that could have been derived from the original dataset) encompass a vast and complex area [9] and are living a new golden age after the publication of the generative adversarial networks (GANs) where one neural network compete to generate new synthetic data that can fool a discriminator network [7]. Even further, the recent success of stable diffusion [24], and other models that can generate detailed images from text descriptions, are pushing the known limits of the generative networks field. These advancements forecast a unexplored landscape of possibilities for innovative applications and improvements.

Autoencoders are a well-known neural network model from the generative model family successfully applied in image processing especially to reconstruct images. They are mainly used in different fields for nonlinear dimensionality reduction and automatic feature learning [16]. Combined with convolutional neural networks (CNN), they can extract features from images in an unsupervised manner, preventing the descriptors bias introduced by subjective decisions. This approach has proven useful to learn complex features in a large variety of fields where images have a prominent role, such as animation [19], neurobiology [22], and medical imagery [26]. In agriculture, images are increasingly used for plant and crop phenotyping [17, 12], providing relevant information to undertake genomic analysis. For example, [30] applied DL methods in images obtained by mobile vehicles to extract high-throughput phenotypic information have served to study the genetic architecture of flowering time in wheat. Such a strategy is particularly useful for traits that are difficult to describe by other means. It is worth mentioning that most of the important agronomic traits in crops are complex and vaguely described by simple descriptors, for they can be dissected into a complex combination of characteristics. This is the case of fruit quality, which embraces visual (fruit color, shape, symmetry), organoleptic (taste, flavor, texture), and sensory (firmness) aspects. Therefore, acquiring relevant data challenges genomic studies. In the case of traits related to fruit or plant appearance, the direct use of images could defeat this difficulty [25]. Since the translation of such images into measurable relevant traits challenges their use in genomic studies, feed whole images to CNN models appears as a good opportunity. Here we present an autoencoder framework to predict fruit shape from known molecular data (SNPs) retrieving, as output, the predicted fruit shape as an image. This model could be successfully used to predict other traits in other species or plant organs.

## 2 Materials and methods

### 2.1 Data

#### 2.1.1 Apple images acquisition and processing

In this study we used the apple images generated in Dujak et al. 2023. Shortly, the images contained from six to ten apple halves from 356 genotypes of the apple reference collection (RefPop) [14] grown at the IRTA experimental field of Gimenells (Lleida, Spain). The apples were collected in 2019 (247 genotypes) and 2020 (the 247 genotypes of 2019 plus 109) at ripening stage. Single apple images were segmented using a fine-tuned version of Mask-RCNN [10] into individual images of size 300×300 RGB pixels (Figure S 3). Apples were classified into five categories depending on their external fruit shape index (FSI): flat, flat-globose, globose, oval and oblong [15].

#### 2.1.2 Genotypic data

A matrix of 303,239 SNP data for the 356 apple genotypes was retrieved from the apple RefPop [14] and prepared for the machine learning process by encoding the dataset to a numerical scale (0, 0.5, 1 representing AA, Aa, aa alleles, respectively).

To train the predictor model, two SNP datasets were extracted from the matrix: one of 150 size and shape-related SNPs (tSNPs) from literature [13] or identified by GWAS in our laboratory (data not published), and the one of 150 randomly selected SNPs (rSNPs) using NumPy library [8]. The random selection of SNPs was renewed every time a new model was trained.

### 2.2 Neural networks models: autoencoder and embedding predictor

The neural networks were developed using PyTorch [20]. First, a convolutional autoencoder network [6] was created, which reduced the number of the image parameters while keeping the relevant features related to the edges and shapes of the apple sections. The architecture used for this convolutional autoencoder network consisted of six convolutions with kernel size of 3×3, set the stride to two and used a rectified linear activation function (ReLu). The number of filters were progressively increased (from 16 to 128) through the multiple encoder layers. The architecture finished with three linear layers with a leaky ReLU activation and sizes of 8192, 4096, and 2048 and with a final linear layer of 64 units with sigmoid activation (Figure 1, Encoder). The decoder followed the inverse structure with a final sigmoid activation in the last layer(Figure 1, Decoder). The model was compiled using the perceptual loss of the VGG network [33] as the loss function, and the stochastic gradient descent (SGD [23]) algorithm was used as optimizer. Training was performed for 35 epochs, over apple cuts of the training dataset.

**Figure 1:**
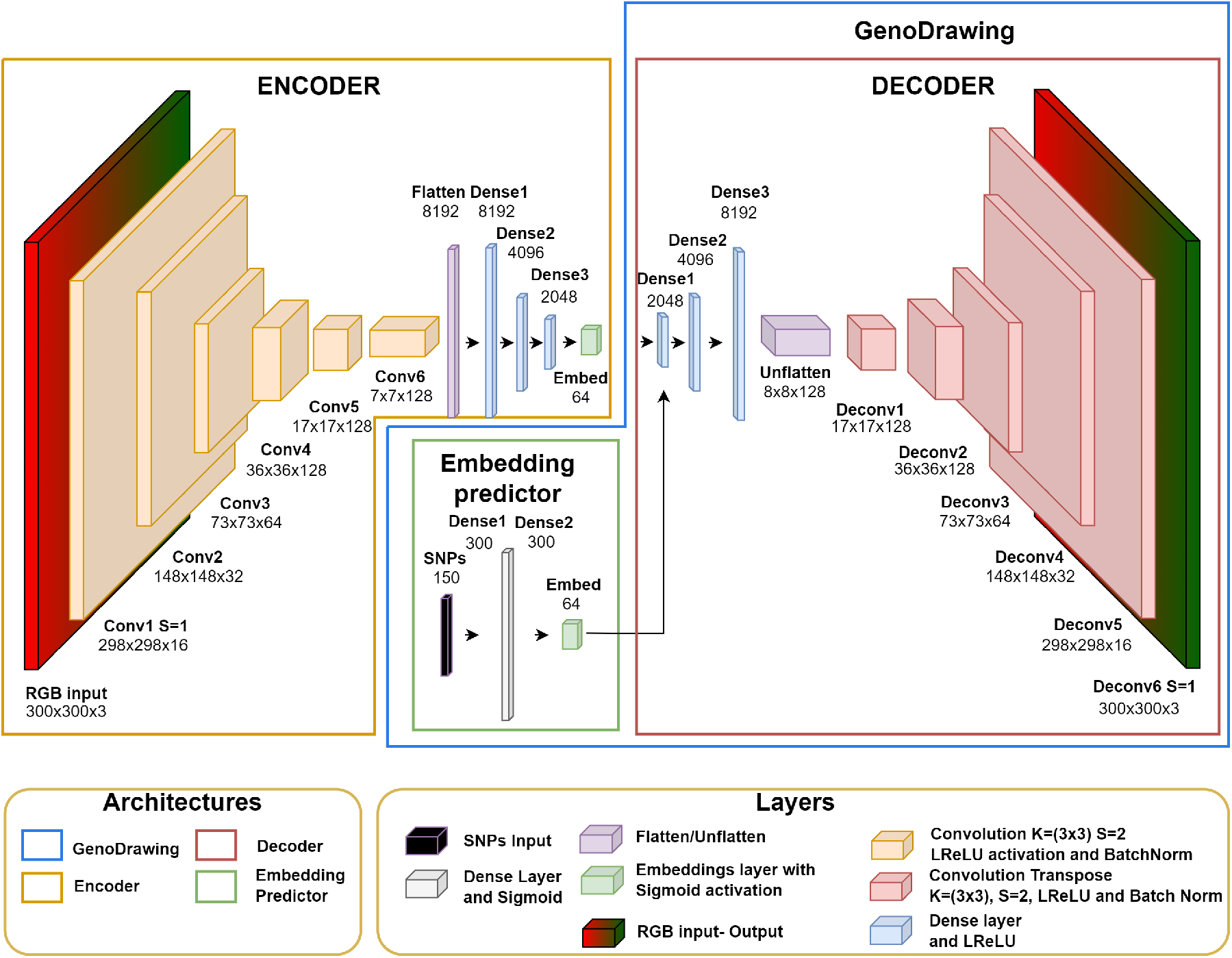
Neural networks architecture schema. The autoencoder is split into encoder and decoder. The encoder, orange box, is compound of six convolutions with a three final dense layers to encode the image into twelve embeddings. The decoder, red box, uses three dense layers and six deconvolutions to recover the image from the embeddings. The embedding predictor, green box, estimates the embeddings values for every genotype. Together, embedding predictor and encoder, compose GenoDrawing, blue box.

The SNPs and embedding values of each genotype in the training dataset were used to train a two layers model to generate an embedding predictor model. This model consisted of i) a linear layer of 300 units with sigmoid activation, which received as input the 150 SNPs; and ii) a 64 units linear layer with sigmoid activation which connected with the 300 units layer and served later as a connection to the decoder when the model was fully trained (Figure 1, Embedding predictor). The embedding predictor was then compiled using mean absolute error (MAE) as the loss function and the SGD algorithm as optimizer (learning rate of 0.05 with scheduler to reduce on plateau) and then trained for 1000 epochs with an early stopper watching for over-fitting symptoms. The resulting model was attached to the decoder part of the autoencoder model generating a SNPs-to-Image model, the GenoDrawing (Figure 1, GenoDrawing). No end-to-end training was performed at this stage.

### 2.3 Embedding predictor target dataset generation

To generate a target dataset for the embedding predictor model, we utilized a trained autoencoder to reduce the dimensionality of each image to a vector of 64 values. Subsequently, each vector representing an image was grouped by genotype, and a mean vector was computed. This mean vector represented the target for the embedding predictor models and was also utilized in the decoder to produce a mean image representing the mean image for the genotype (Figure 2). The resulting dataset of generated images was utilized in the evaluation of the metrics by estimating their FSI and their SR.

**Figure 2:**
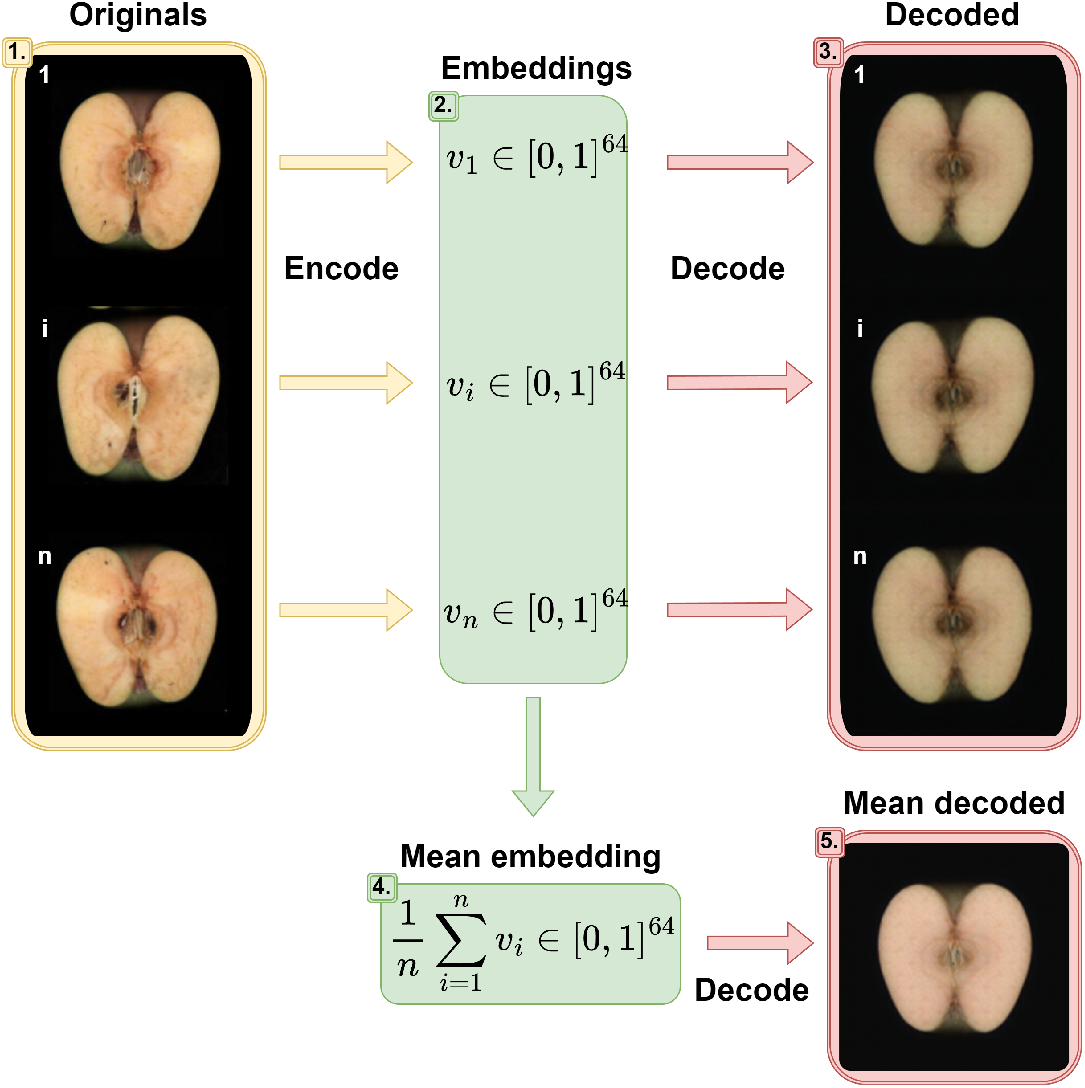
The original images (1) were encoded into vectors of size 64 (2) using the encoder. These vectors could be decoded into images (3). Also, the vectors were used to estimate a mean embedding value per genotype (4). The mean embeddings were used to produce a mean image per genotype (5). These mean embedding values were used to train the embedding predictor, and the mean images were used to evaluate the resulting GenoDrawing.

## 3 Results

### 3.1 General approach

The general approach (Figure 3) consisted of training two models: 1) an image compressor model (autoencoder), trained with images of apple sections, and 2) an embedding predictor, a model that predicts the mean compressed values, also known as embeddings, of a genotype based on its single nucleotide polymorphisms (SNPs). Both models were then merged into a SNPs-to-Image model which we named GenoDrawing.

**Figure 3:**
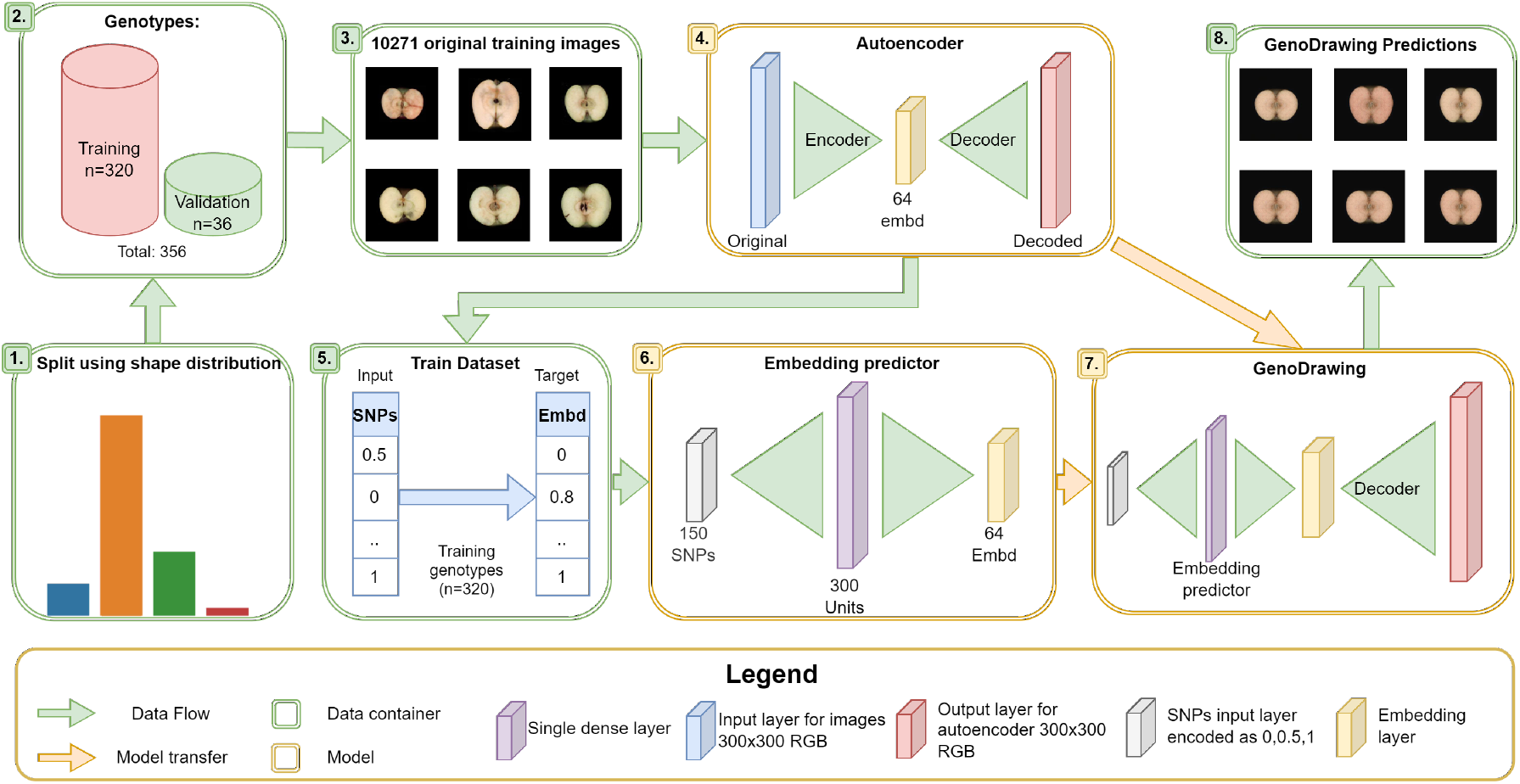
Graphical abstract of the general approach. The dataset was divided into two splits conserving each similar fruit shape distribution (1,2). Images from the training dataset were used to fit the autoencoder (3,4). Encoded values for each image (with 64 embeddings) were averaged by genotype. and used together with SNPs to train the embedding predictor (5,6). GenoDrawing is not a trained model but an assembly of both, autoencoders decoder, and embedding predictor (7). Images predicted by GenoDrawing(8).

Images were separated into a training and a validation datasets conserving similar shape typologies distributions (Figure 3.1). The training dataset contained the 90% of the images (10271 of 320 genotypes). The rest (1340 of 36 genotypes) were used for validation (Figure 3.2). The autoencoder model was fitted with the training dataset to ultimately produce vectors of 64 embedding values per image (Figure 3.3-4). These vectors were used to generate a 10271×64 matrix, which was then reduced to a 320×64 matrix using the mean embedding values per genotype. The autoencoder model was then used in the 320×64 matrix to retrieve the average image for each genotype. The same process was followed for the validation dataset without retraining the image compressor.

Image and genotypic datasets (Figure 3.5) were used for training and validating the embedding predictor model. The dataset of random SNPs was renewed in each iteration (Figure 3.6). The decoder section of the autoencoder and the embedding predictor were assembled into the GenoDrawing to retrieve the predicted image (Figure 3.7-8).

### 3.2 Autoencoder and embedding predictor training

To reduce the complexity of the apple images we used the autoencoder network, which was able to compress (encode) each into 64 dimensions (embeddings), number that was selected by trial and error through multiple tests with the number of embeddings ranging from 12 to 1024. The reconstructed (decoded) images from the 64 embeddings maintained, in a great measure, the shape aspect of the original image while the flesh color and lobule depth were not reproduced (Figure 4). The selected dimensionality allowed for the preservation of important information in a small dimensional space, which served as the target for prediction. During the autoencoder training, the perceptual loss, which indicates the resemblance of the reconstructed image to the original, improved consistently from around 0.15 to a 0.09 along the 35 epochs. The mean encoder values per genotype were used together with SNP data to train the embedding predictor and reconstruct the apple section images. The embedding predictor was trained through the number of epochs needed until an early stopper was activated. For all the SNP datasets tested, a slow decrement in the mean absolute error was observed through the various iterations of training on the validation dataset, with a stagnation that arrived earlier or later depending on the number of SNPs used. There was no evidence of overfitting.

**Figure 4:**
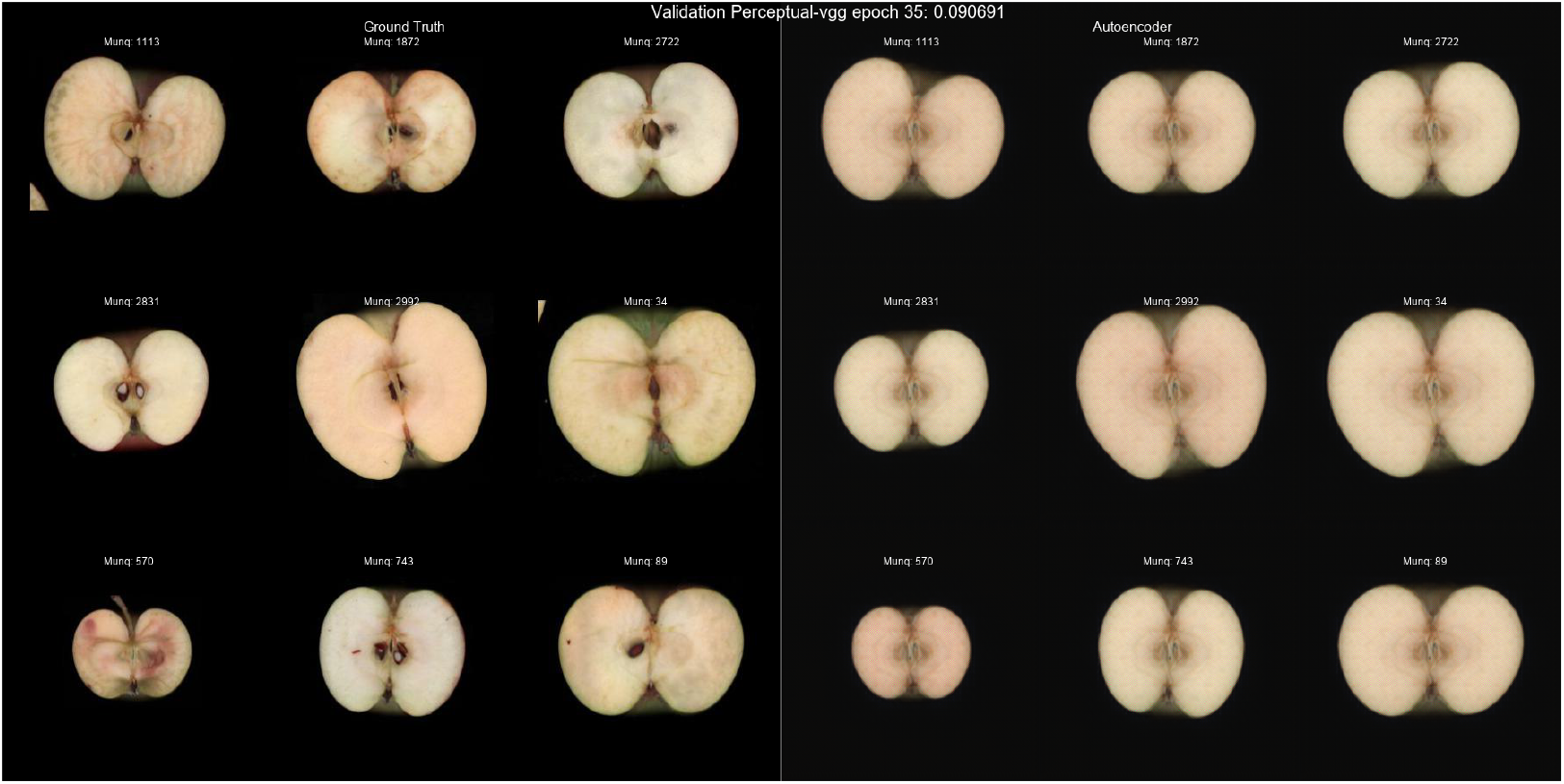
On the left side, original apple section. On the right side, output given by the autoencoder for the correspondent apple.

### 3.3 Comparing the impact of trait-related and randomly selected SNPs on embedding prediction models for GenoDrawing

To compare the accuracy of the embededing prediction model using targeted or random SNPs, we trained 100 models for each case. Using tSNPs consistently led to lower mean absolute error (MAE) values. On average, the MAE values obtained using tSNPs was 0.0864 with a standard deviation of 0.0034. When rSNPs were used, the average MAE was 0.0944 with a standard deviation of 0.0017 (Figure S1).

The best two embedding predictors among the 100 previously described (MAE 0.0857 and 0.0889 for the targeted and random SNP datasets, respectively) were used along with the decoder model to generate the predicted images and, eventually, test the accuracy of GenoDrawing. Mean decoded and predicted images were analyzed to determine the Fruit Shape Index External (FSI [11]), and the Shoulder Ratio (SR, ratio between the width at the most top and bottom points of the apple)(Figure S3). Deviation of FSI and SR values with the tSNP dataset was lower, with MAE of 0.0386 and 0.0317 for the FSI and SR, respectively, compared to MAE of 0.0651 and 0.0493 for the same to metrics in the rSNPs dataset. To determine the quality threshold for the MAE values, the mean and standard deviation of the FSI and SR were used to produce 10,000 datasets of 36 samples. These datasets were generated according to a *N*(*μ,σ^2^)* distribution based on the validation dataset, and subsequently compared to the FSI and SR of the images produced by GenoDrawing. The resulting MAE superior boundary was established at 0.075 with a standard deviation of 0.0084 for the FSI, and at 0.0529 with a standard deviation of 0.0054 for the SR. Given that these boundary values exceed those produced by both rSNPs and tSNPs derived models, it may be concluded that the predictions are not merely the result of random sampling from the original distribution.

Once the non-randomness of the models had been confirmed, the distributions corresponding to both the rSNPs and tSNPs based versions of the embedding prediction were compared to the decoded means and the original data distributions for each genotype in the validation dataset (Figure 5). From Wasserstein distances between the distributions we infer that tSNPs models generate distributions closer to the ones of the Decoded and Genotype Means, 0.0209 and 0.0259 respectively, than the rSNPs models, (0.0469 and 0.0331, respectively) (Table S1). Additionally, it was observed that there exists a small difference between the distribution of the decoded mean image and the original images. Specifically, the Wasserstein distance values between the decoded mean distribution and the original image distributions were relatively small for both the FSI attribute, 0.0144, and the SR attribute, 0.01748.

**Figure 5:**
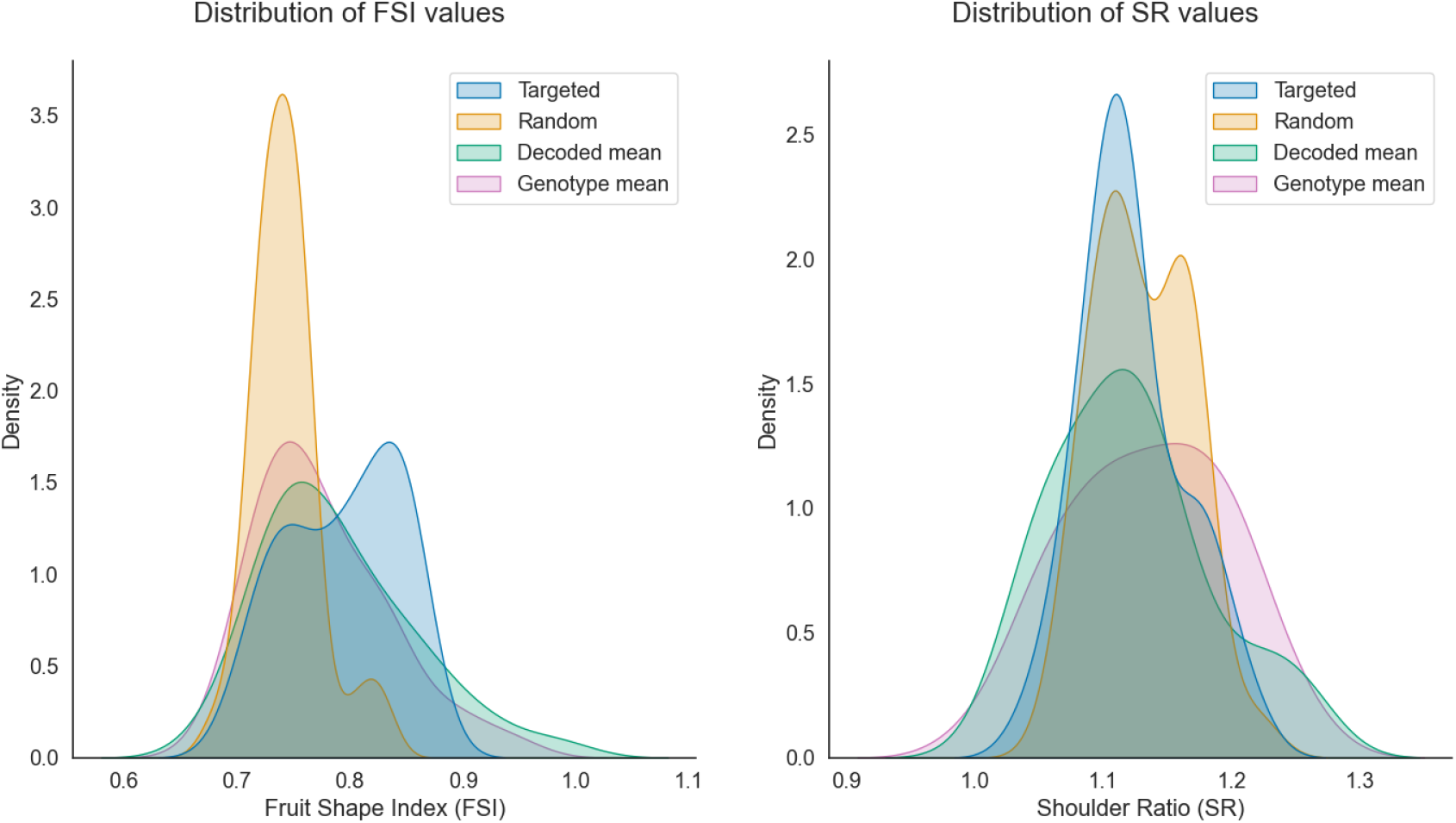
The distribution plots for two different traits: Fruit Shape Index (FSI) on the left and Shoulder Ratio (SR) on the right. The distributions were generated using the GenoDrawing approach with the best-scoring version of the embedding predictor for both trait-related SNPs (represented in blue) and random SNPs (represented in orange). For comparison, the distributions of the validation dataset were also plotted using the mean images (represented in green) and the original images (represented in red).

To further understand the limitations of the proposed model, the autoencoder and predicted images were classified, based on the FSI, into five categories (flat, flat globose, globose, oblong, and oval) [15]. The success in class assignation was assessed using the accuracy and F-Score metrics and plotted in a confussion matrix (Figure S4). Both metrics were consistently lower for the tSNPs based version with an accuracy of 0.61 and F-score of 0.59 against an accuracy of 0.39 and F-Score of 0.34 for the random version. Furthermore the categories provided by the rSNPs based version only comprehended two categories, belonging most of the predictions to the flat category, while the targeted version was able to produce in three categories with a better distribution. None of them was able to produce images belonging to the oblong and oval categories.

### 3.4 Effect of Adding Non-Shape-Related Markers on Embedding Predictor Model Accuracy

To investigate the effect of increasing the number of markers selected without knowing the biological relevance, we added to the tSNP dataset an increasing number of SNPs in nine steps up to a total of: 150, 300, 1650, 4650, 10150, 20150, 50150, 100150, and 300150. We used a similar approach with the random-selected SNPs, increasing the size of the selection at each step. For each of these steps and datasets, 15 models were trained. The embedding predictor models with only 150 tSNPs were generally more effective in achieving a low mean absolute error (MAE) compared to models that include additional SNPs. The exception was observed when using 300150 targeted version, where three outliers showed lower MAEs than their random selection counterparts. Notably, a unique case was observed where a targeted version consisting of 300 SNPs achieved the lowest observed MAE of 0.0835 (Figure 6).

**Figure 6:**
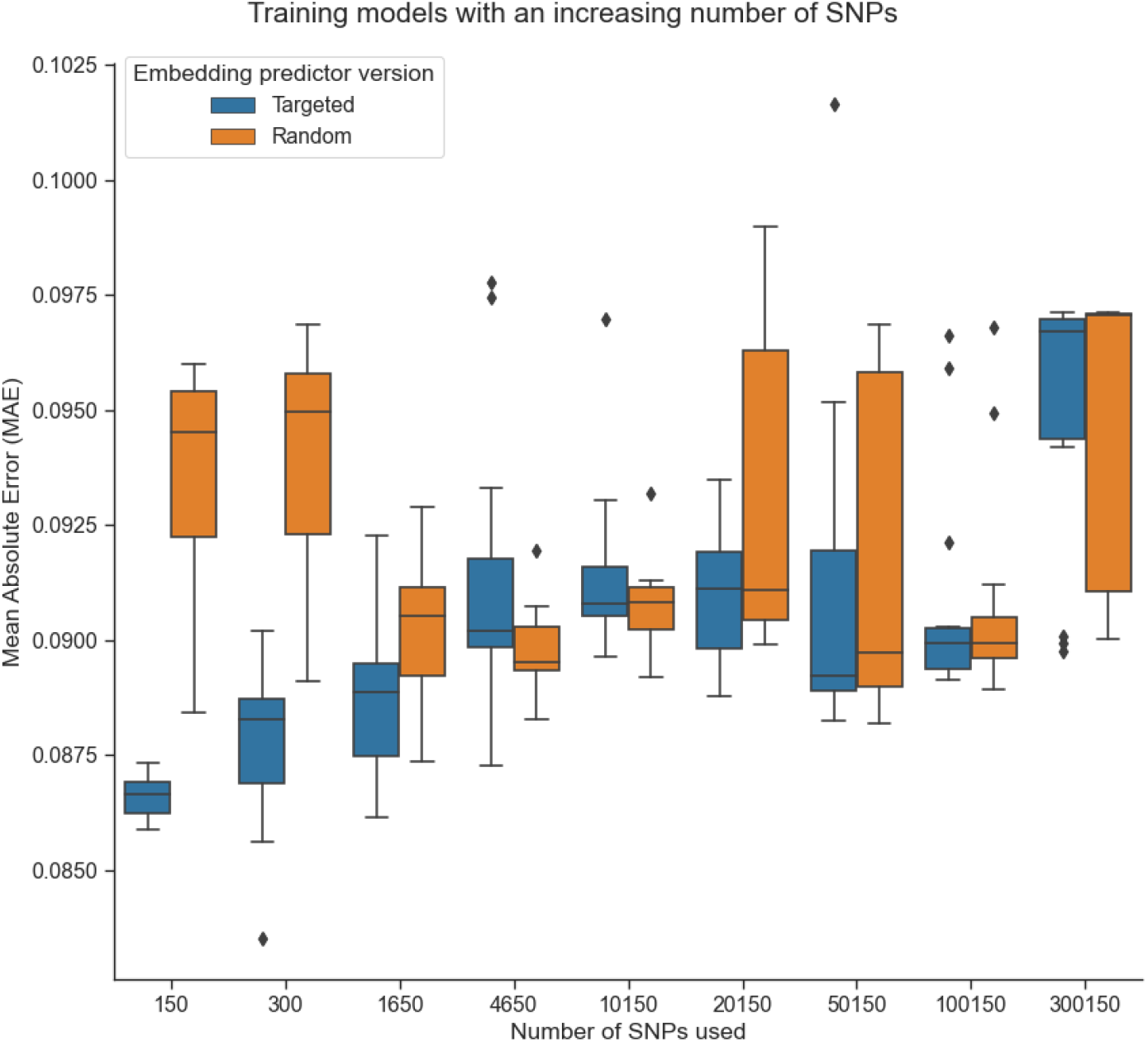
Embedding predictor models with increasing size. Two versions of the models were observed: a targeted version that always included 150 SNPs relevant to shape as well as random markers, denoted by the color blue, and a random version which involved a fully random selection of markers, denoted by the color orange. Fifteen models were trained for each of the different sizes and versions.

## 4 Discussion

### 4.1 Evaluation of Autoencoder Performance, Loss Function Selection, and Embedding Predictor Structure

In this study, we employed an autoencoder network to simplify the complexity of the apple images into 64 embeddings. This number of embeddings was deemed sufficient to account for small variations in the images while circumventing the complexity associated with adjusting a large number of embeddings to a reduced dataset of SNPs. Although changes in the number of embeddings can produce different outcomes, given the scope of this study, we selected the number that aligned better with our objectives. The images produced from the embeddings successfully captured a reliable and straightforward representation of the apples, effectively reproducing the original image structure. Nevertheless, certain features, such as lobule depth and flesh color, were not accurately represented, likely due to the sparsity of the embedding space and the difficulty faced by a simple autoencoder network in generating photorealistic images. As the primary focus of this study was on shape, the autoencoder structure was deemed appropriate for the task. However, future studies that aim to capture a wider range of attributes may benefit from utilizing a different structure to better accomplish the reconstruction task.

With respect to the loss function utilized in our study, we evaluated the Mean Square Error (MSE), SSIM-based metrics, and perceptual loss. We decided on the perceptual loss given the well proved adequacy to the reconstruction task [33] and the results provided. Although SSIM-based metrics, such as MS-SSIM and CW-SSIM, are robust in terms of comparing image structures, they often overlook many details [27]. In our experiments the reconstructions were similar to the original images in terms of shape but were less visually alike. Furthermore, MSE tendency to generate blurry boundaries between figures [31] proved to be highly detrimental to the comparison of image shapes in our study, preventing a proper examination of the data. this

The embedding predictor structure employed in this study was relatively simple, consisting of two single layers with 300 units each and sigmoid activation. This approach was selected following an examination of several architectures, including deeper models with skip connections, convolutions, and direct models, on the 150 SNP version predictions. The simplicity of this approach allowed for the exploration of different input sizes while minimizing training time. For larger datasets, however, more complex architectures capable of studying sequences while retaining position relevance could prove beneficial. Nevertheless, the primary goal of this study was to assess the feasibility of this approach and the impact of selecting the appropriate SNPs relevant to the features studied.

### 4.2 Using the mean embedding as a representation of the phenotype

One of the primary challenges associated with utilizing an autoencoder is determining which information is being encoded in the embedding space. In our study, a relevant question that arose was whether the mean of the embeddings represented the mean of the images corresponding to a specific genotype. To evaluate this, we compared the distribution of both for the FSI and SR metrics. We found out that the FSI distributions were highly similar both in distributions and images; however, the shoulder ratio differed significantly (Figure 5). This was probably due to the asymmetry of the images. An asymmetric apple that is cut in two halves will produce two asymmetric mirror images, which will be then merged in a symmetric mean image. This issue could be tackled in the future by removing from the dataset one of the halves of each apple.

### 4.3 Relevance of SNP selection in genome prediction models

Genome prediction models are typically founded on dense SNP data. Nevertheless, generating and managing large SNP datasets remains a challenge for small to medium-sized breeding companies, despite the relatively low cost of high-throughput genotyping methods. Strategies such as genotyping-by-sequencing (GBS) have been developed to reduce genotyping costs [4]; however, the use of smaller arrays would be more conducive to the implementation of this methodology. To this end, we reduced the genotyping array to a small array of 150 SNPs that we found to be significant for different shape traits. We tested the model with both random and selected SNPs and consistently observed an improvement in results when using the selected SNPs. This supports the notion that the selected SNPs are indeed significant and might represent a good health sign for their relationship with apple shape (Figure S1). However, we also identified a rare case in which the addition of random SNPs to the selected SNPs led to better results, leading us to believe that our selection may be missing some shape-relevant SNPs. It is clear that the inclusion of relevant SNPs is crucial for accurate predictions, which is consistent with the biological interpretation of SNP information. In theory, the presence of the relevant SNPs should be sufficient for making predictions, regardless of the addition of random SNPs. Our results demonstrate a consistent improvement in performance when the relevant SNPs are included in the input, with a decrease in performance as the input size increases. This decrease in performance could be attributed to the simplicity of the embedding prediction structure used in our study. As previously mentioned, a more complex architecture may be necessary for larger inputs to better extract relevant information, which can be addressed through architecture improvements.

### 4.4 Evaluating embedding predictor performance using FSI and SR metrics in GenoDrawing

Two metrics were employed to evaluate the similarity in fruit shape between the mean image generated by encoding all images for a particular genotype and their prediction. The Fruit Shape Index (FSI) was used as a reliable measure of the general shape of the fruit, while the shoulder ratio was used to capture the conicity of the apple. However, it was noted that the shoulder ratio was not adequately captured by using the mean embedding, limiting the interpretability of the results obtained, but still providing valuable insight into the challenges of using this model to predict asymmetric metrics. Conversely, the FSI metric was found to be well-suited to the task. The results, as shown in Figure 5, revealed that the random embedding predictor was highly focused on a limited range of variability for predictions. This indicated that the embedding predictor consistently produced similar images to minimize the error, but did not effectively learn to solve the task. In contrast, the targeted version produced a better-distributed range of predictions, which could achieve higher FSI values and result in better approximations of the image production task. This was further supported by the categorization of predicted images into five shape categories (Figure S4) using FSI scores, which revealed that the random version only predicted in two categories (flat and flat-globose), while the targeted version predicted in three categories (from flat to globose). Overall, these findings suggest that GenoDrawing can learn the task effectively when relevant markers for shape are provided.

### 4.5 Limitations

By estimating the mean shape appearance for a genotype we aimed to cover the variability within genotype, but apples can be very influenced by environmental effect, leading to a wide range of phenotypes within the same individual [5]. Even though most of the genotypes counted with multiple biological replicates and two year imaging, this is one of the biggest challenges faced in genomic prediction. Furthermore, the overrepresentation of flat-globose apples limits the capacity of the model trained, forcing it to be biased toward this category (Figure S2). Removing the number of examples from this category has been considered, but this considerably would limit the size of the dataset in term of number of genotypes which might hurt the learning process more than helping it. For traits with such a low separation between classes a larger genotype dataset, including more extreme phenotypes may help to improve accuracy in future studies.

## 5 Conclusions

Through the GenoDrawing SNP-to-Image method presented in this work, we propose a new framework to study genomic prediction for fruit trees phenotype. Our study showed that using trait-associated SNPs significantly improves the performance of GenoDrawing in predicting fruit shape compared to using random SNP selections. The consistent better optimization during training with trait-associated SNPs and the fact that the results are within the dataset distribution boundaries also provide evidence that this methodology is moving in the right direction. Our approach of using a low number of SNPs to generate predictions at a relatively low cost might be particularly relevant for small to medium breeding companies. Additionally, the same approach could be applied to predict other heritable traits with higher economic impact such as physiological disorders or genetic disease severity. We propose GenoDrawing method as a feasible framework for studying genomic prediction for phenotyping in fruit trees, which could lead to less biased categorizations for shape, color, and other parameters that are currently predicted through subjective categories. The use of specific genetic knowledge within the problem can significantly improve the accuracy of the model, as observed in our study.

**Table 1:** Wasserstein distance between distributions based on the Fruit Shape Index (FSI) and Shoulder Ratio (SR) for four different cases: Targeted, Random, Decoded Mean, and Genotype Mean. The diagonal shows the distance between identical distributions, and the upper/lower triangles shows the distances between different distributions.

|  |  |  |  |  |
| --- | --- | --- | --- | --- |
| **FSI** | Targeted | Random | Decoded mean | Genotype mean |
| Targeted | 0 | 0.0505 | 0.0209 | 0.0259 |
| Random |  | 0 | 0.0469 | 0.0331 |
| Decoded mean |  |  | 0 | 0.0144 |
| Genotype mean |  |  |  | 0 |
| **SR** | Targeted | Random | Decoded mean | Genotype mean |
| Targeted | 0 | 0.0108 | 0.0184 | 0.0255 |
| Random |  | 0 | 0.0247 | 0.0233 |
| decoded mean |  |  | 0 | 0.0175 |
| Genotype mean |  |  |  | 0 |

## 6 Acknowledgments

FJR is recipient of grant PRE2019-087427 funded by MCIN/AEI/ 10.13039/501100011033 and by “ESF Investing in your future. This research was supported by project PID2021-128885OB-I00 funded by MCIN/AEI/10.13039/501100011033 and by “ERDF A way of making Europe”. This project has received funding from the European Union’s Horizon 2020 research and innovation programme under grant agreement No 817970 (INVITE). We acknowledge support from the CERCA Programme (“Generalitat de Catalunya”), and the “Severo Ochoa Programme for Centres of Excellence in R&D” 2016-2019 (SEV-2015-0533) and 2020-2023 (CEX2019-000902-S) both funded by MCIN/AEI /10.13039/501100011033.

## 7 Author contributions

The work was conceived and designed by FJR and MJA. DR, JAB provided intellectual support. FJR and MJA wrote the manuscript. All authors revised the manuscript.

## 8 Data availability

The code repository including notebooks, and models with their trained weights can be found in the following github repository: https://github.com/Fedjurrui/GenoDrawing

The image and SNP datasets will be shared under request.

## 9 Supplementary Material

**Figure S 1:**
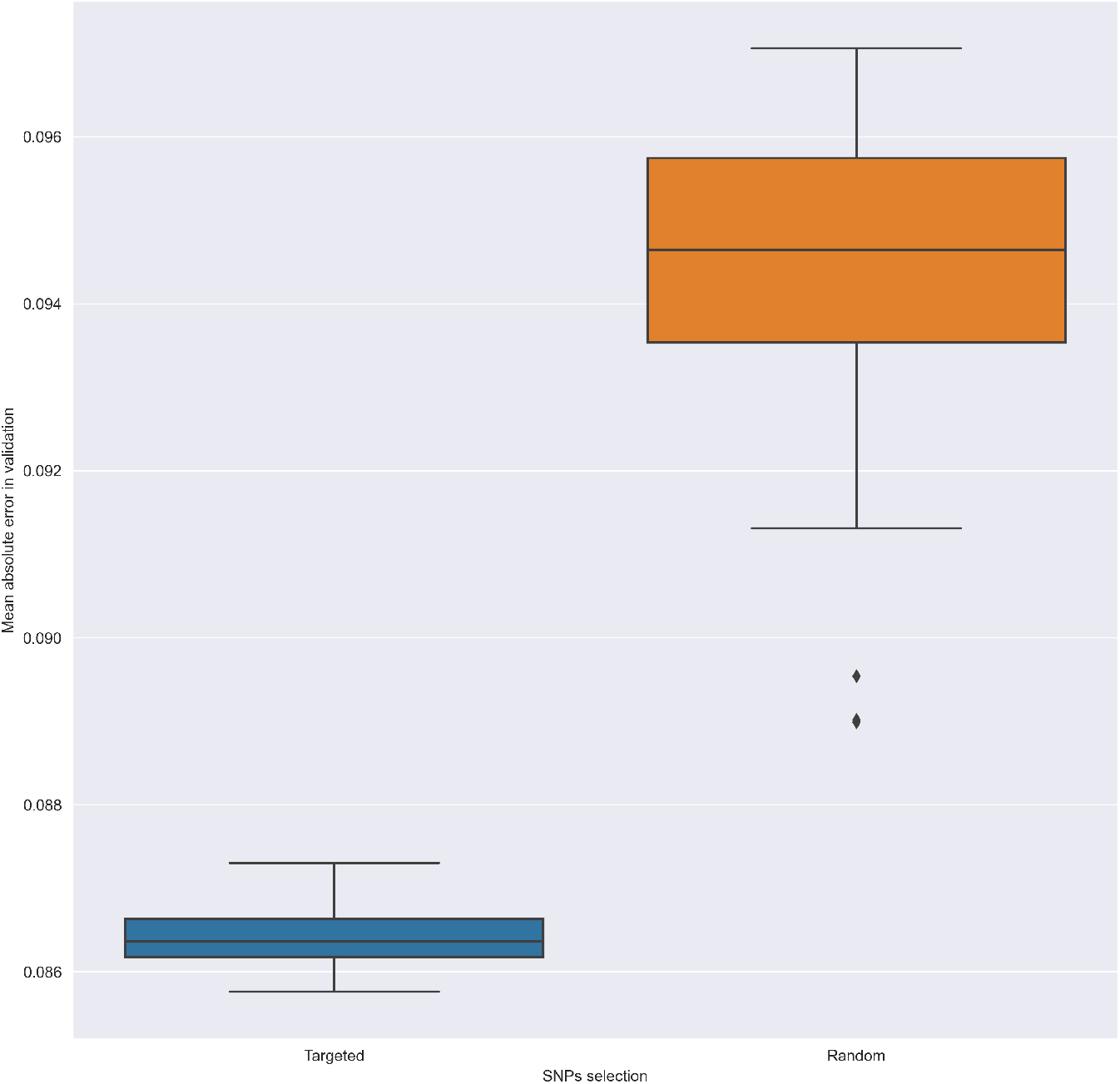
Training Losses for both versions of the SNP to embedding predictor. The training was performed one hundred times for each.

**Figure S 2:**
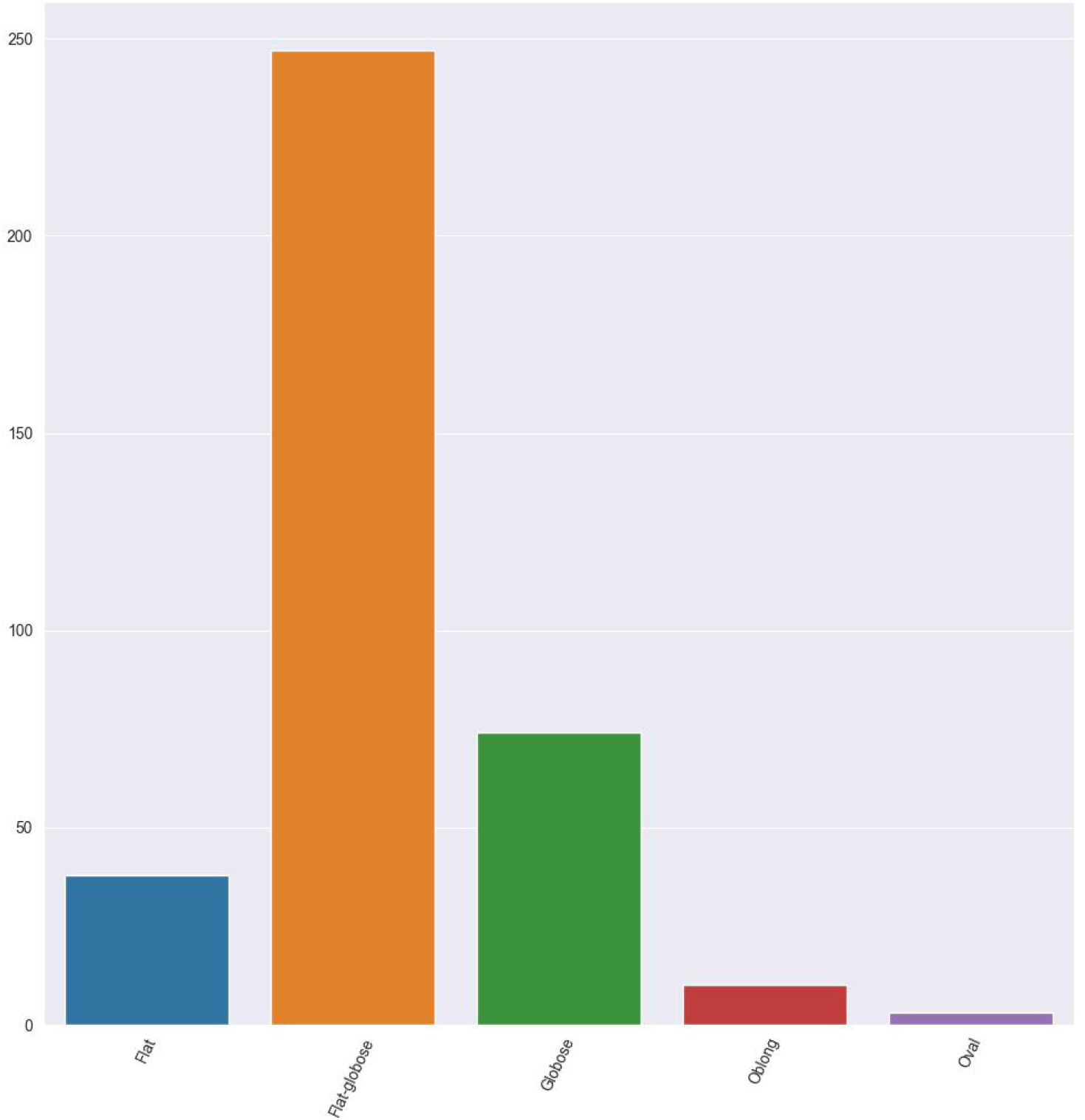
Dataset shape categories distribution.

**Figure S 3:**
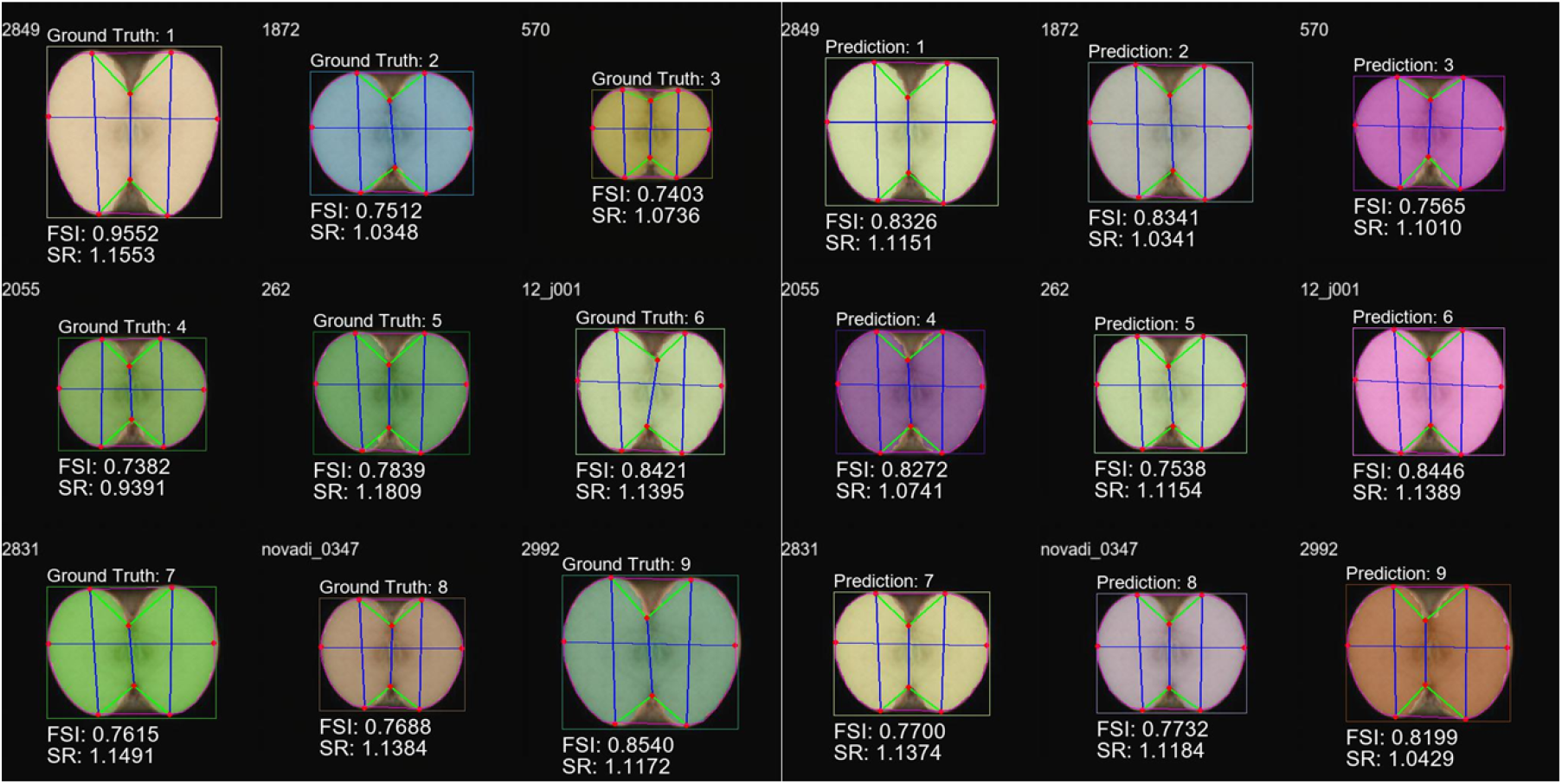
Left, nine examples produced through the mean embedding values for the genotype in the validation dataset. Right, the nine corresponding predictions for the GenoDrawing model with relevant-to-shape SNPs. The inner contour detection is displayed in colors; the red lines are the measures used to calculate to form Fruit Shape Index External (FSI) and Shoulder ratios (SR).

**Figure S 4:**
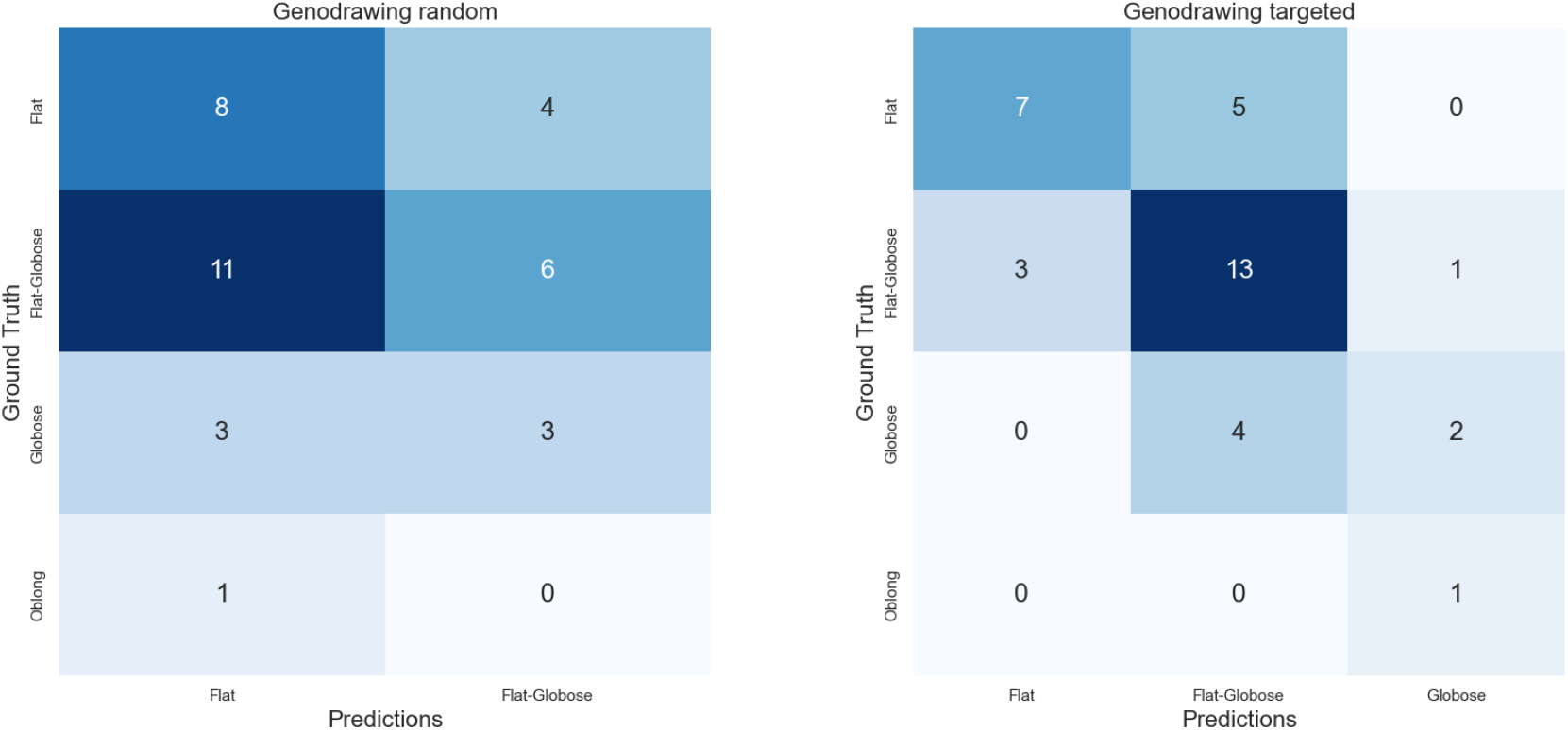
Left, the confusion matrix for the classification in ECPGR categories using the predictions generated by the model with a random selection of genomic markers. Right the same classification but using the model with a selection of genomic markers based on a shape GWAS

## References

[1] Martin Abadi, Ashish Agarwal, Paul Barham, Eugene Brevdo, Zhifeng Chen, Craig Citro, Greg S. Corrado, Andy Davis, Jeffrey Dean, Matthieu Devin, Sanjay Ghemawat, Ian Goodfellow, Andrew Harp, Geoffrey Irving, Michael Isard, Yangqing Jia, Rafal Jozefowicz, Lukasz Kaiser, Manjunath Kudlur, Josh Levenberg, Dandelion Mané, Rajat Monga, Sherry Moore, Derek Murray, Chris Olah, Mike Schuster, Jonathon Shlens, Benoit Steiner, Ilya Sutskever, Kunal Talwar, Paul Tucker, Vincent Vanhoucke, Vijay Vasudevan, Fernanda Viégas, Oriol Vinyals, Pete Warden, Martin Wattenberg, Martin Wicke, Yuan Yu, and Xiaoqiang Zheng. Tensorflow: Large-scale machine learning on heterogeneous systems, 2015. Software available from tensorflow.org.

[2] Rostam Abdollahi-Arpanahi, Daniel Gianola, and Francisco Peñagaricano. Deep learning versus parametric and ensemble methods for genomic prediction of complex phenotypes. Genetics Selection Evolution, 52(1):12, Feb 2020.

[3] Javaid A. Bhat, Sajad Ali, Romesh K. Salgotra, Zahoor A. Mir, Sutapa Dutta, Vasudha Jadon, Anshika Tyagi, Muntazir Mushtaq, Neelu Jain, Pradeep K. Singh, and et al. Genomic selection in the era of next generation sequencing for complex traits in plant breeding. Frontiers in Genetics, 7, 2016.

[4] Robert J. Elshire, Jeffrey C. Glaubitz, Qi Sun, Jesse A. Poland, Ken Kawamoto, Edward S. Buckler, and Sharon E. Mitchell. A robust, simple genotyping-by-sequencing (gbs) approach for high diversity species. PLOS ONE, 6(5):1–10, 05 2011.

[5] Toshi M Foster and Maria José Aranzana. Attention sports fans! The far-reaching contributions of bud sport mutants to horticulture and plant biology. Horticulture Research, 5, 07 2018. 44.

[6] Ian Goodfellow, Yoshua Bengio, and Aaron Courville. Deep learning, 2016. http://www.deeplearningbook.org.

[7] Ian Goodfellow, Jean Pouget-Abadie, Mehdi Mirza, Bing Xu, David Warde-Farley, Sherjil Ozair, Aaron Courville, and Yoshua Bengio. Generative adversarial networks. COMMUNICATIONS OF THE ACM, 63, 2020.

[8] Charles R. Harris, K. Jarrod Millman, Stéfan J. van der Walt, Ralf Gommers, Pauli Virtanen, David Cournapeau, Eric Wieser, Julian Taylor, Sebastian Berg, Nathaniel J. Smith, Robert Kern, Matti Picus, Stephan Hoyer, Marten H. van Kerkwijk, Matthew Brett, Allan Haldane, Jaime Fernández del Río, Mark Wiebe, Pearu Peterson, Pierre Gérard-Marchant, Kevin Sheppard, Tyler Reddy, Warren Weckesser, Hameer Abbasi, Christoph Gohlke, and Travis E. Oliphant. Array programming with NumPy. Nature, 585(7825):357–362, September 2020.

[9] GM Harshvardhan, Mahendra Kumar Gourisaria, Manjusha Pandey, and Siddharth Swarup Rautaray. A comprehensive survey and analysis of generative models in machine learning. Computer Science Review, 38:100285, 2020.

[10] Kaiming He, Georgia Gkioxari, Piotr Dollár, and Ross Girshick. Mask r-cnn. In 2017 IEEE International Conference on Computer Vision (ICCV), pages 2980–2988, 2017.

[11] Huan Huan, Yong Sun, Chun Bo Zhao and, Dong Min Li and, Mei Chen, Xin Yi Wang and, Zhen Zhong Zhang and, and Hai Han. Identification of markers linked to major gene loci involved in determination of fruit shape index of apples (malus domestica).

[12] Sumit Jangra, Vrantika Chaudhary, Ram C Yadav, and Neelam R Yadav. High-throughput phenotyping: A platform to accelerate crop improvement. Phenomics, 1:31–53, 2021.

[13] Michaela Jung, Beat Keller, Morgane Roth, Maria José Aranzana, Annemarie Auwerkerken, Walter Guerra, Mehdi Al-Rifaï, Mariusz Lewandowski, Nadia Sanin, Marijn Rymenants, Frédérique Didelot, Christian Dujak, Carolina Font i Forcada, Andrea Knauf, François Laurens, Bruno Studer, Hélène Muranty, and Andrea Patocchi. Genetic architecture and genomic predictive ability of apple quantitative traits across environments. Horticulture Research, 9, 02 2022. uhac028.

[14] Michaela Jung, Morgane Roth, Maria José Aranzana, Annemarie Auwerkerken, Marco Bink, Caroline Denancé, Christian Dujak, Charles-Eric Durel, Carolina Font i Forcada, Celia M Cantin, Walter Guerra, Nicholas P Howard, Beat Keller, Mariusz Lewandowski, Matthew Ordidge, Marijn Rymenants, Nadia Sanin, Bruno Studer, Edward Zurawicz, Françcois Laurens, Andrea Patocchi, and Hélène Muranty. The apple REFPOP—a reference population for genomics-assisted breeding in apple. Horticulture Research, 7, 11 2020. 189.

[15] Marc Lateur, E Dapena, David Szalatnay, A Guyader, Inger Hjalmarsson, Monika Höfer, M Militaru, Carlos Miranda Jiménez, Gregor Osterc, Alain Rondia, et al. ECPGR Characterization and Evaluation Descriptors for Apple Genetic Resources: Apple (Malus X Domestica). ECPGR-European Cooperative Programme for Plant Genetic Resources, 2022.

[16] Walter Hugo Lopez Pinaya, Sandra Vieira, Rafael Garcia-Dias, and Andrea Mechelli. Chapter 11 - autoencoders. In Andrea Mechelli and Sandra Vieira, editors, Machine Learning, pages 193–208. Academic Press, 2020.

[17] Keiichi Mochida, Satoru Koda, Komaki Inoue, Takashi Hirayama, Shojiro Tanaka, Ryuei Nishii, and Farid Melgani. Computer vision-based phenotyping for improvement of plant productivity: a machine learning perspective. 8:1–12, 2018.

[18] Osval A. Montesinos-López, Abelardo Montesinos-López, Roberto Tuberosa, Marco Maccaferri, Giuseppe Sciara, Karim Ammar, and José Crossa. Multi-trait, multi-environment genomic prediction of durum wheat with genomic best linear unbiased predictor and deep learning methods. Frontiers in Plant Science, 10, 2019.

[19] Lucas Mourot, Ludovic Hoyet, François Le Clerc, François Schnitzler, and Pierre Hellier. A survey on deep learning for skeleton-based human animation. Computer Graphics Forum, 41(1):122–157, 2022.

[20] Adam Paszke, Sam Gross, Francisco Massa, Adam Lerer, James Bradbury, Gregory Chanan, Trevor Killeen, Zeming Lin, Natalia Gimelshein, Luca Antiga, Alban Desmaison, Andreas Kopf, Edward Yang, Zachary DeVito, Martin Raison, Alykhan Tejani, Sasank Chilamkurthy, Benoit Steiner, Lu Fang, Junjie Bai, and Soumith Chintala. Pytorch: An imperative style, high-performance deep learning library, 2019.

[21] Miguel Pérez-Enciso and Laura M. Zingaretti. A guide on deep learning for complex trait genomic prediction. Genes, 10(7), 2019.

[22] Blake A. Richards, Timothy P. Lillicrap, Philippe Beaudoin, Yoshua Bengio, Rafal Bogacz, Amelia Christensen, Claudia Clopath, Rui Ponte Costa, Archy de Berker, Surya Ganguli, Colleen J. Gillon, Danijar Hafner, Adam Kepecs, Nikolaus Kriegeskorte, Peter Latham, Grace W. Lindsay, Kenneth D. Miller, Richard Naud, Christopher C. Pack, Panayiota Poirazi, Pieter Roelfsema, João Sacramento, Andrew Saxe, Benjamin Scellier, Anna C. Schapiro, Walter Senn, Greg Wayne, Daniel Yamins, Friedemann Zenke, Joel Zylberberg, Denis Therien, and Konrad P. Kording. A deep learning framework for neuroscience. Nature Neuroscience, 22(11):1761–1770, Nov 2019.

[23] Herbert Robbins and Sutton Monro. A stochastic approximation method. The Annals of Mathematical Statistics, 22(3):400–407, September 1951.

[24] Robin Rombach, Andreas Blattmann, Dominik Lorenz, Patrick Esser, and Björn Ommer. High-resolution image synthesis with latent diffusion models, 2021.

[25] Rijad Sarić, Viet D. Nguyen, Timothy Burge, Oliver Berkowitz, Martin Trtílek, James Whelan, Mathew G. Lewsey, and Edhem Čustovió. Applications of hyperspectral imaging in plant phenotyping. Trends in Plant Science, 27(3):301–315, 2022.

[26] Ying Song, Shuangjia Zheng, Liang Li, Xiang Zhang, Xiaodong Zhang, Ziwang Huang, Jianwen Chen, Ruixuan Wang, Huiying Zhao, Yutian Chong, Jun Shen, Yunfei Zha, and Yuedong Yang. Deep learning enables accurate diagnosis of novel coronavirus (covid-19) with ct images. IEEE/ACM Transactions on Computational Biology and Bioinformatics, 18(6):2775–2780, 2021.

[27] Dogancan Temel and Ghassan AlRegib. Image quality assessment and color difference. In 2014 IEEE Global Conference on Signal and Information Processing (GlobalSIP), pages 970–974, 2014.

[28] Thomas van Klompenburg, Ayalew Kassahun, and Cagatay Catal. Crop yield prediction using machine learning: A systematic literature review. Computers and Electronics in Agriculture, 177:105709, 2020.

[29] Athanasios Voulodimos, Nikolaos Doulamis, Anastasios Doulamis, and Eftychios Protopa-padakis. Deep learning for computer vision: A brief review. Computational Intelligence and Neuroscience, 2018:7068349, Feb 2018.

[30] Xu Wang, Hong Xuan, Byron Evers, Sandesh Shrestha, Robert Pless, and Jesse Poland. High-throughput phenotyping with deep learning gives insight into the genetic architecture of flowering time in wheat. GigaScience, 8(11), 11 2019. giz120.

[31] Zhou Wang and Alan C. Bovik. Mean squared error: Love it or leave it? a new look at signal fidelity measures. IEEE Signal Processing Magazine, 26(1):98–117, 2009.

[32] Haohao Zhang, Lilin Yin, Meiyue Wang, Xiaohui Yuan, and Xiaolei Liu. Factors affecting the accuracy of genomic selection for agricultural economic traits in maize, cattle, and pig populations. Frontiers in Genetics, 10, 2019.

[33] R. Zhang, P. Isola, A. A. Efros, E. Shechtman, and O. Wang. The unreasonable effectiveness of deep features as a perceptual metric. In 2018 IEEE/CVF Conference on Computer Vision and Pattern Recognition (CVPR), pages 586–595, Los Alamitos, CA, USA, jun 2018. IEEE Computer Society.

[34] Y. Zhao, M. F. Mette, M. Gowda, C. F. H. Longin, and J. C. Reif. Bridging the gap between marker-assisted and genomic selection of heading time and plant height in hybrid wheat. Heredity, 112(6):638–645, Jun 2014.

